# GLP-1R is downregulated in beta cells of NOD mice and T1D patients

**DOI:** 10.1101/845776

**Authors:** Asha Recino, Kerry Barkan, Anja Schmidt-Christensen, Julia Nilsson, Nick Holmes, Duncan Howie, Dan Holmberg, Pär Larsson, Malin Flodström-Tullberg, Luca Laraia, David R Spring, Jacob Hecksher-Sørensen, Anne Cooke, Graham Ladds, Maja Wållberg

## Abstract

Glucagon-like peptide 1 (GLP-1) is produced by L cells in the small intestine in response to ingested glucose and increases insulin release from pancreatic beta cells by activation of its cognate receptor (GLP-1R). Stimulation of this receptor also contributes to increased beta cell survival and regeneration. We have found that pancreatic beta cells from Non Obese Diabetic (NOD) mice express significantly lower levels of GLP-1R than C57BL/6 mice, leaving the NOD beta cells with an impaired response to GLP-1 stimulation. The lower expression appears to be caused by accelerated degradation of GLP-1R in the beta cells, a process that can be reversed by inhibiting trafficking to the lysosome. Importantly, our results appear to translate to the human disease since we also observed significantly lower expression of the GLP-1R in pancreatic islets from donors with type 1 diabetes. These results suggest that beta cell physiology may play a role in susceptibility to autoimmune inflammation.

## Introduction

Type 1 diabetes is believed to be caused by immune mediated destruction of the insulin producing beta cells in the pancreas. When these cells are destroyed or incapacitated, the body can no longer effectively take up glucose, leading to glucose starved tissues combined with excess levels of glucose in the blood (1, 2).

Despite decades of research, the event or events that precipitate disease remain unknown. It is clear that certain genes, especially within the MHC locus which directs T cell specificity, predispose to type 1 diabetes (3). Scientists have justifiably focussed on the immune response in efforts to understand the disease and to find a cure. However, there is increasing interest in how much the beta cell contributes to its own destruction, and whether beta cell fragility may be a contributing factor in disease susceptibility (4-6). More than half of the known genes associated with type 1 diabetes are expressed in the beta cell, and their expression increases when the cell is stressed (7). Beta cells can be stressed through different mechanisms including prolonged exposure to high levels of glucose and exposure to proinflammatory cytokines and other chemicals (8). It is therefore possible that abnormal and/or prolonged stress responses predispose to type 1 diabetes through turning the beta cell into a more available and vulnerable target. The presence of residual beta cells in pancreas tissue from deceased patients with long standing disease (9) also indicates that the lack of adequate insulin responses in patients could be due to dysfunction of these cells in addition to destruction.

Beta cells produce insulin in response to elevated levels of glucose. This effect can be potentiated by the hormone glucagon-like peptide 1 (GLP-1) (10) which is produced by cells in the small intestine in response to ingested glucose (11, 12). Due to the actions of GLP-1, beta cells secrete increased amounts of insulin in response to ingested glucose when compared to the same concentration of glucose injected intravenously (13), a phenomenon described as the incretin effect(14). GLP-1 is short lived *in vivo* due to rapid inactivation by dipeptidylpeptidase-4 (DPP-4)(11), but inhibitors of DPP-4 as well as the DPP-4-resistant GLP-1 analogue exendin-4 are used successfully to increase insulin release in type 2 diabetes patients (15). As stimulation of the GLP-1 receptor (GLP-1R) has been reported to protect beta cells and even induce regeneration of the beta cell pool (16, 17), attempts have been made to combine these effects with other therapies to prevent or treat type 1 diabetes in mice and humans (18-22). These studies have shown either no, or very limited, effects of GLP-1R stimulation in reversing or preventing type 1 diabetes.

We have found that non-obese diabetic (NOD) mouse islets have impaired responses to GLP-1 as well as exendin-4. This impairment is due to a global reduction in GLP-1R in the beta cell that appears to be caused by accelerated degradation. Such lowered expression, and therefore impaired response to GLP-1, render NOD beta cells functionally impaired which may increase their susceptibility to autoimmune attack. We also observe lower expression of GLP-1R on beta cells of human type 1 diabetic patients. Our findings provide a possible explanation for the limited success of GLP-1 analogues in treating type 1 diabetes.

## Results

### NOD mice manifest deficient responses to GLP-1 analogue exendin-4

GLP-1 and its analogues stimulate release of insulin from beta cells, which in turn allows uptake of glucose from the blood into tissues. Wildtype C57BL/6 mice respond to injection of the GLP-1 analogue exendin-4 by a drop in the levels of blood glucose (Fig. 1a). We found that NOD mice do not respond to exendin-4 injections by the expected drop of blood glucose concentration (Fig. 1a). This is not just an effect of the autoimmune attack on beta cells in diabetic NOD mice, as the same effect can be observed in B- and T cell deficient NOD^*scid*^ mice in which the beta cells are completely unharmed by inflammation (Fig. 1a). The decreased effect of exendin-4 is not specific to the NOD mouse colony at the Department of Pathology, as this was also observed in other NOD mouse colonies (Supplementary figure 1a, 1b), and the effect does not appear to be dominant, as F1 offspring of a C57BL/6xNOD cross do exhibit a response to injection of exendin-4 (Fig. 1a). Further, using exendin-4 labelled with Cy3, we observed binding to beta cells in the islets of C57BL/6 mice(23) (Fig. 1b, middle right panel), but importantly only a very weak binding occurred on the islets of the NOD mice (Fig. 1b, middle left panel). In addition, the labelled exendin-4 could be easily detected in the kidneys in both NOD^*scid*^ and C57BL/6 mice, where it is excreted (Fig. 1b, bottom panels). Islets in both NOD^*scid*^ and C57BL/6 mice stained well for insulin and glucagon, indicating that they were functional and otherwise appeared healthy (Fig. 1b, top panels). Finally, we noted that by increasing the dose to 4 nmol per mouse (ca 8×10^5^ fold higher than the physiological levels of GLP-1) we could observe a reduction in blood glucose, albeit attenuated, in the NOD^*scid*^ mice, indicating that the NOD mice do express some receptor (Supplementary figure 1c, d) but that they require much higher concentrations of ligand to achieve even a modest response.

**Figure 1.**
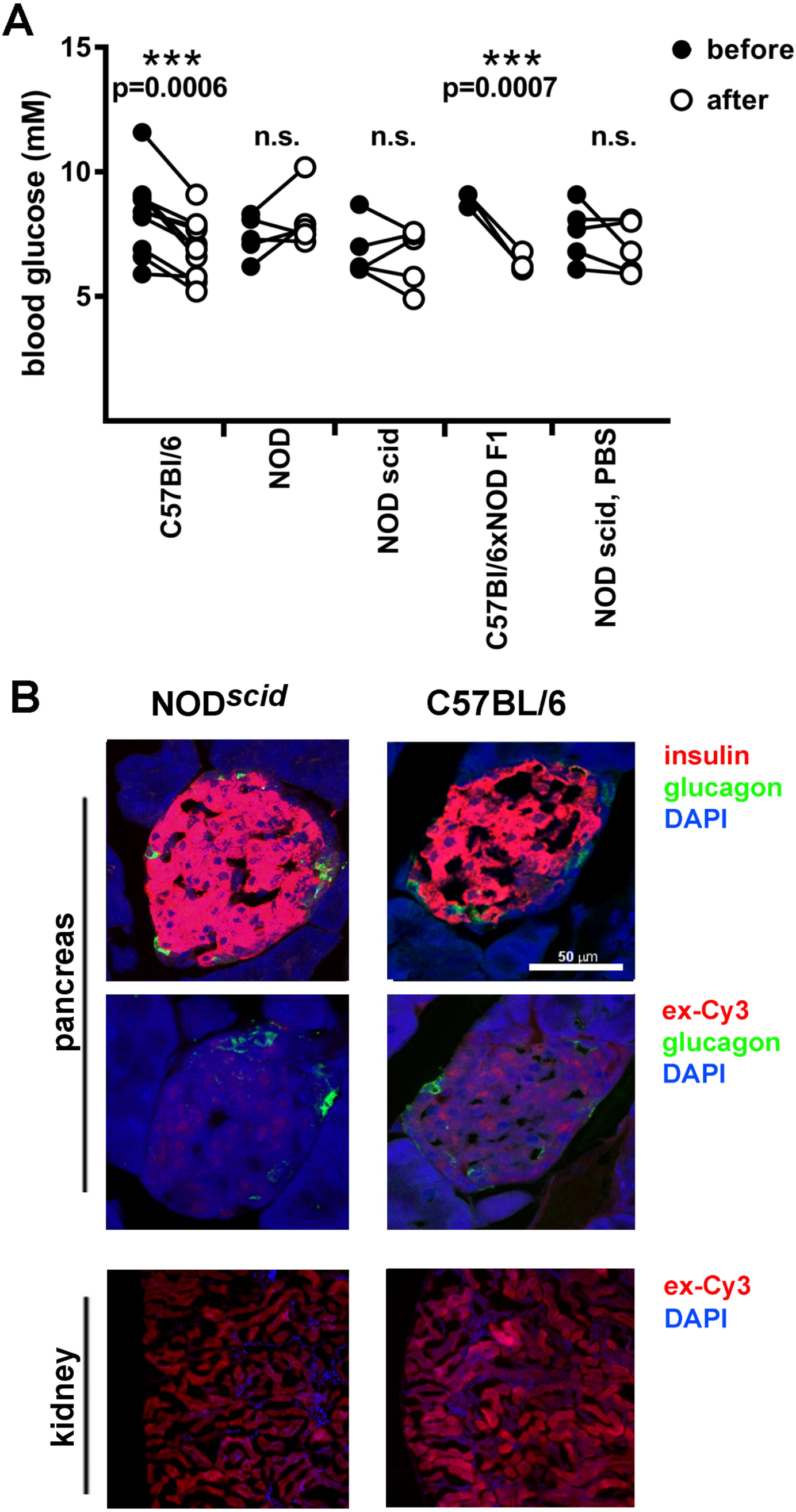
GLP-1 analogue exendin-4 does not bind or activate islets in mice on a NOD genetic background. (A) Measurements of blood glucose were taken before and 1 h after injection of 400 pmol exendin-4 in C57BL/6, NOD, NOD^*scid*^ and C57BL/6 x NOD F1 mice, or an equal volume of vehicle (PBS) in NOD^*scid*^ mice. Differences were measured using a paired t-test comparing before and after blood glucose levels. Each dot represents one mouse, and the experiment was repeated at least three times except for the F1 cross which was performed once. (B) Binding of Cy3-labelled exendin-4 (red) (4nmol iv) was assessed in islets from either NOD^*scid*^ mice (middle left panel) and C57BL/6 (middle right panel), with glucagon (green) indicating the location of alpha cells. Insulin stain (B, top panels) was equal in both strains, and labelled exendin-4 could be detected in the kidney, where it is excreted, in both strains (B, bottom panels).

### GLP-1 receptor gene expression is normal in NOD mouse islets, and the distribution of the GLP-1 analogue exendin-4 in other organs is unchanged

We wondered if the lack of activity and binding of exendin-4 could be due to a mutation in the gene encoding for GLP-1R. However, investigation of the NOD and C57BL/6 *glp-1r* sequence from the Sanger web site (24) showed no differences in the sequence between the strains. Islets of NOD^*scid*^ mice expressed GLP-1R mRNA at similar levels to C57BL/6 islets (Fig. 2a). We detected the highest levels of GLP-1R mRNA in picked islets and whole pancreas and lower but still detectable levels in lung and the pyloric valve of the stomach. GLP-1R expression has been reported in brain (25, 26), heart (27, 28), kidney (29) and aorta (26), but the levels (where given) are at least 100-fold lower than in beta cells and in some cases observed in highly enriched sub-fractions (e.g. brain). We do not rule out that there are levels of mRNA present which are too low to be significant relative to the housekeeping reference, or diluted by the use of unfractionated tissue. There were no differences in expression between the mouse strains in any of the organs (Fig. 2a). We therefore asked if the lack of binding and responses in *in vivo* injected mice could be due to excessive expression of receptor in other tissues, which would sequester the exendin-4. By injecting Cy3-labelled exendin-4 and examining different organs one hour post injection we observed similar biodistribution of the labelled ligand in lung and stomach, while the difference in fluorescence in the pancreas was consistent with our previous results (Fig. 2b). The labelled exendin-4 peptide was excreted via the kidney, hence the very high signal in this organ.

**Figure 2.**
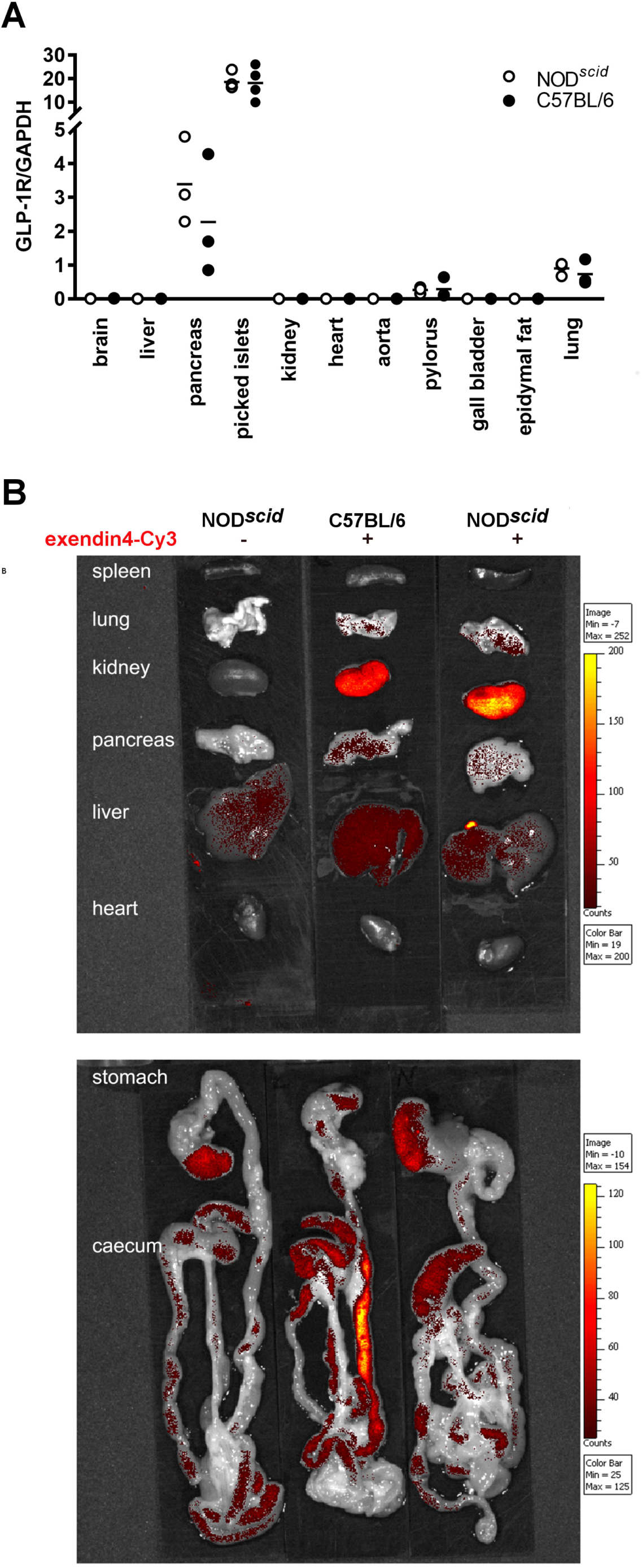
GLP-1 receptor gene expression is normal in NOD^*scid*^ mouse islets, and the distribution of Cy3-labelled exendin-4 in other organs is unchanged. (A) TaqMan PCR analysis of expression of *glp1r* in tissues collected from NOD^*scid*^ and C57BL/6 was assessed using *gapdh* as a housekeeping control. (B) Accumulation of Cy3-labelled exendin-4 in different organs of either NOD^*scid*^ (B, right panels) and C57BL/6 mice (B, middle panels) was assessed using IVIS imaging. A vehicle-injected control was used to determine background signal (B, left panels).

### Deficient binding of Cy3-labelled exendin-4 depends on properties of the NOD^*scid*^ islet rather than external factors

To test whether the lack of binding of exendin-4 to NOD islets *in vivo* is due to properties of the NOD islet itself or the environment within which it resides, such as soluble factors or binding to other tissue, we performed cross-transplantation experiments. We transplanted NOD^*scid*^ or C57BL/6 islets under the kidney capsule of either NOD^*scid*^ or C57BL/6 recipients, and then investigated whether the transplanted islets could bind injected labelled exendin4 probe. We found that the islets retained the exendin-4 binding properties of the donor even after transplantation, so that C57BL/6 islets stained with the injected probe regardless of whether the recipient was NOD^*scid*^ or C57BL/6, while NOD^*scid*^ islets did not (Fig. 3a, bottom panels). An insulin stain was performed on adjacent sections to ascertain that the islets were healthy (Fig. 3a, top panels). To support this data, we assessed *in vivo* binding of the labelled peptide to islets by real-time noninvasive 2-Photon imaging. As described before, the exendin-4 peptide acts as an agonist on the GLP-1R, thus not only binding it, but also inducing increased insulin release. As the mice had to be anaesthetised for the *in vivo* imaging procedure and we did not want to risk inducing hypoglycaemia, we used a non-agonistic exendin-9 peptide, which binds GLP-1R but does not initialise signalling and is not internalised (30). Pancreatic islets of noninflamed NOD^*Rag2-/-*^ or C57BL/6 mice were transplanted into the anterior chamber of the eye (ACE) of immunodeficient C57BL/6^*Rag2-/-*^ or NOD^*Rag2-/-*^ mice and repeatedly imaged (Fig. 3b, left panel) (31, 32). After an initial phase of vascularization, allogenic islets transplanted into the ACE of immunodeficient mice are sustained and functional as demonstrated by reflection (Supplementary figure 2). Finally, injection of NOD^*Rag2-/-*^ and C57BL/6^*Rag2-/-*^ recipient mice with labelled exendin-9 peptide showed that although binding to the NOD^*Rag2-/-*^ islets could be observed, it was considerably dimmer than in the C57BL/6 islets (Figure 3b, right and middle panel).

**Figure 3.**
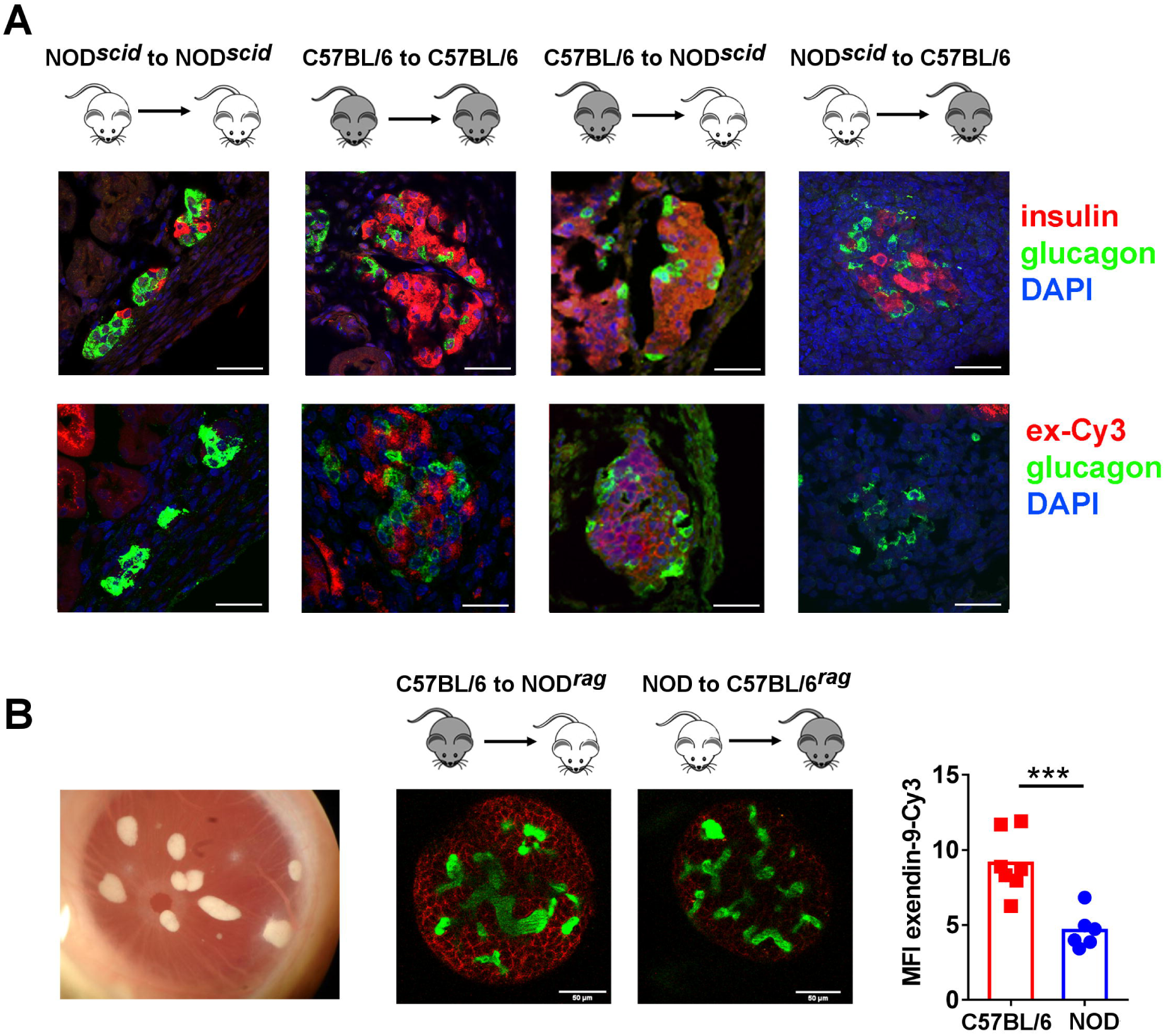
Deficient binding of Cy3-labelled exendin-4 depends on properties of the NOD^*scid*^ islet rather than external factors. (A) Islets were isolated from male NOD^*scid*^ or C57BL/6 mice (10-12 weeks old) and transplanted under the kidney capsule of either male NOD^*scid*^ or C57BL/6 recipients as indicated in the figure. After 10 days, the mice were given an injection of 4 nmol (ca 2μg) of exendin-4-Cy3 iv. 1 hour after the injection, the graft bearing kidney was harvested stained for insulin and glucagon (A, top panels) or glucagon alone to counter stain for the exendin-4 labelling (A, bottom panels). (B) Islets were transplanted into the anterior chamber of the eye (ACE) (B, left panel). Islets from albino C57BL/6 mice were transplanted into the ACE of a NOD^*Rag2-/-*^ recipient mice (B, middle left panel) and islets from NOD^*Rag2-/-*^ mice were transplanted into albino C57BL/6^*Rag2-/-*^ mice (B, middle right panel). 10 weeks post-transplant, the mice received an injection of 2 nmol Cy3-labelled exendin-9 (red), and binding was assessed *in vivo* using 2Photon imaging. Blood vessels are visualised by iv injection of dextran-FITC in green. Evaluation of staining intensity in different transplanted islets is plotted in B, right panel. The difference between the groups was assessed using the Student’s t-test.

### NOD^*scid*^ islets express less GLP-1R and are hyporesponsive to GLP-1R stimulation *in vitro*

Previous work has demonstrated that labelled exendin-4 can be used for flow cytometric sorting of NOD beta cells(33), although this work did not comment on relative brightness of the signal. Immunofluorescent staining of pancreas sections showed very weak GLP-1R antibody staining of the NOD^*scid*^ islets (Fig. 4a, right panel), compared to C57BL/6 (Fig. 4a, left panel). This was confirmed with flow cytometry, determining the level of fluorescence of isolated islet cells incubated with saturating levels of Cy3-labelled probe. This showed presence of binding in NOD^*scid*^ beta cells but at a lower intensity than in the C57BL/6 cells (Fig. 4b). The lower expression of GLP-1R was not an indication that all surface receptors are downregulated in NOD^*scid*^ beta cells, as similar levels of the transferrin receptor, which also cycles between the cell membrane and the endosomal compartment, was equally expressed in NOD^*scid*^ beta cells and C57BL/6 beta cells (Supplementary figure 3)

**Figure 4.**
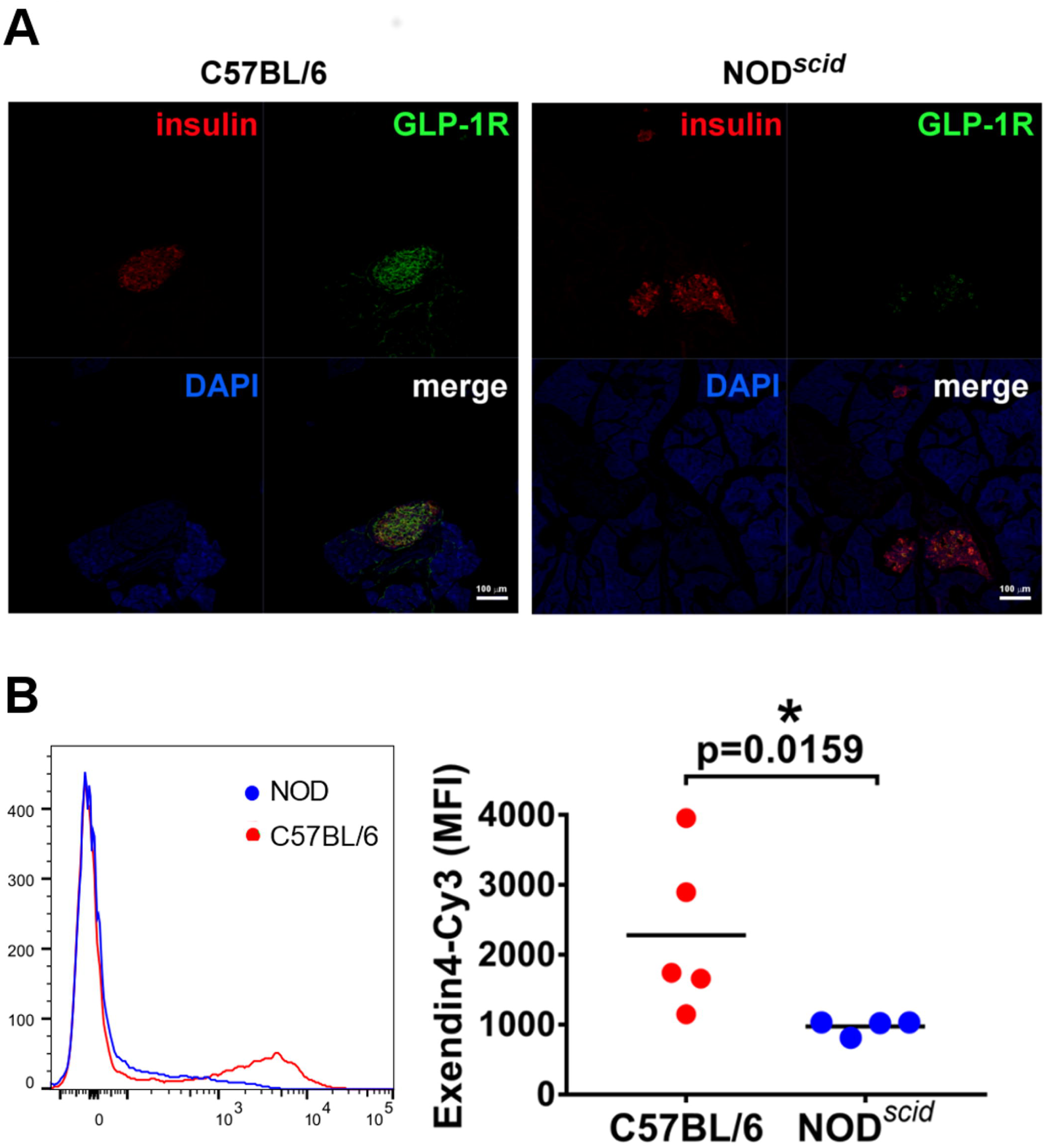
NOD^*scid*^ islets express less GLP-1R. (A) Section staining of GLP-1R in islets in the pancreas using fluorescence microscopy in islets from C57BL/6 (A, left) and NOD^*scid*^ (A, right). (B) Binding of Cy3-labelled exendin-4 probe to beta cells *in vitro* was assessed using flowcytometry. A representative histogram (B, left panel), and a graph plotting all MFI values in the experiment (B, right panel), are shown. Each dot represents one mouse and the difference between the groups was assessed using a Mann-Whitney test (B, right panel).

We next sought to pharmacologically characterise the *in vitro* signalling response (specifically cAMP accumulation) of the NOD^*scid*^ islets when challenged with wild type GLP-1. We observed that the C57BL/6 islets were able to generate cAMP with a significantly higher efficacy (between 10-100 fold increased) compared to the islets isolated from the NOD^*scid*^ islets (Fig. 5a). This data further confirms the reduced functionality of the GLP-1R in the NOD^*scid*^ islets. In addition, insulin secretion was potentiated in an exendin-4 concentration-dependent manner in picked islets from C57BL/6 mice stimulated for 30 minutes with 11mM glucose and the indicated concentrations of exendin-4, while potentiation was attenuated in NOD^*scid*^ islets (Fig. 5b). No Ca^2+^ was added to the medium to minimise non-GLP-1R mediated insulin secretion (34) (Fig. 5b). cAMP signalling after stimulation of the gastric-inhibitory polypeptide receptor (GIPR), another incretin receptor involved in glucose-stimulated insulin release after ingestion of glucose (35), was not suppressed (Fig. 5c).

**Figure 5.**
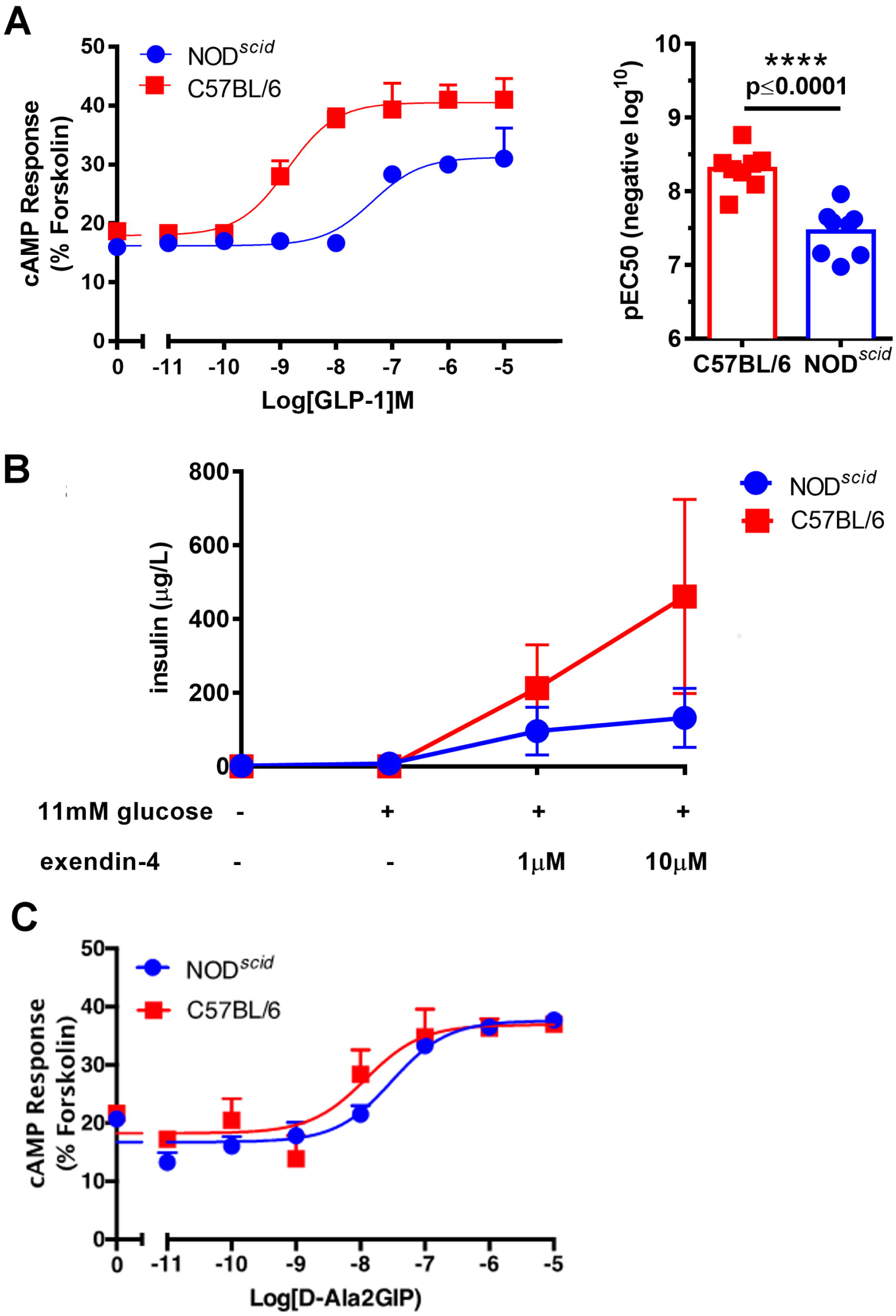
NOD^*scid*^ islets are hyporesponsive to GLP-1 *in vitro.* (A) NOD^*scid*^ and C57BL/6 beta cells were stimulated with GLP-1 *in vitro*, and their cAMP response was determined. The response curve (A, left panel), and the Student’s t-test of the recorded pEC50 (A, right panel) are shown. (B) NOD^*scid*^ and C57BL/6 islets were incubated in 0 or 11mM glucose *in vitro*, in the presence of exendin-4 as indicated. After 30 minutes, supernatants were collected and insulin content was determined. (C) NOD^*scid*^ and C57BL/6 beta cells were stimulated with GIP *in vitro*, and their cAMP response was determined.

### GLP-1R localizes to the lysosome in NOD^*scid*^ beta cells, and protein levels can be restored through inhibition of degradation

As low levels of GLP-1R can be detected in NOD beta cells, we next investigated where in the cell the receptor was enriched. We found no evidence of localisation to the endoplasmic reticulum or the Golgi (Supplementary figure 4), but could detect increased overlap between GLP-1R and the lysosomal compartment through co-staining with lysotracker (Fig. 6a), indicating that the degradation of the receptor is accelerated in NOD beta cells. Interestingly, we found increased levels of lysosomal activity in the NOD cells (Supplementary figure 5). To clarify why the GLP-1R ends up in the lysosome, we investigated whether inhibition of protein degradation could restore GLP-1R expression (Fig. 6b-e). We found that inhibition of lysosome formation through addition of chloroquine could completely restore GLP-1R expression (Fig. 6c and 6e), and that inhibition of autophagy with bafilomycin could partially restore expression (Fig. 6d and 6e). Combination of bafilomycin and chloroquine also resulted in complete restoration of receptor expression, but importantly, insulin levels were not reduced, indicating that, at least in the 4h treatment window, the beta cells integrity and insulin content remains unchanged (Supplementary figure 6). Inhibition of proteasome activity had no effect (data not included). Importantly, pre-incubation of islets with chloroquine before exposure to 11mM glucose with added exendin significantly increased insulin secretion from the NOD islets (Fig. 6f), indicating that chloroquine-mediated rescue of receptor expression also restores GLP-1R signalling effects.

**Figure 6.**
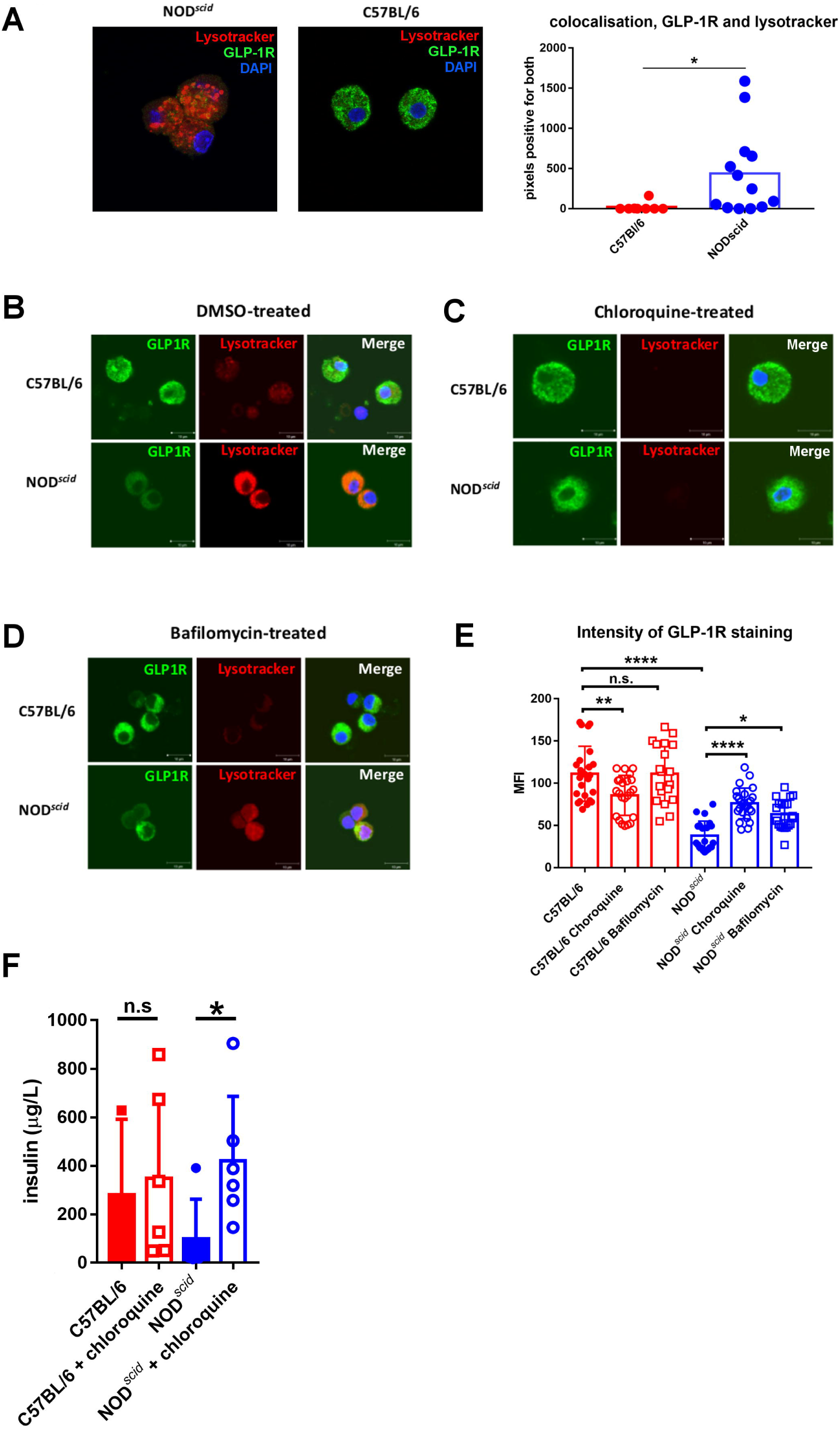
GLP1-R localizes to the lysosome in NOD^*scid*^ beta cells, and protein levels can be restored through inhibition of degradation. (A) NOD^*scid*^ and C57BL/6 beta cells were stained with GLP-1R antibody (green) and lysotracker (red) (A, left and middle panels), and the overlap of the stains was assessed (A, right panel). Each dot represents one cell, and the difference between the groups was determined using the Student’s t-test. Data is representative of three independent experiments. (B-D) GLP-1R, lysotracker and DAPI staining in C57BL/6 (B-D, top panels) and NOD^*scid*^ beta cells (B-D, bottom panels) after incubation with vehicle alone (DMSO) (B), Chloroquine (C) and Bafilomycin (D). (E) Graph representation of GLP-1R mean fluorescence intensity in the beta cells after culture. Differences between groups were assessed using ANOVA followed by a Dunnett’s post test for multiple testing. Each dot represents one cell. (F) Insulin secretion from C57BL/6 islets or NOD^*scid*^ islets pre-incubated with either vehicle (control) or chloroquine (cq), and then stimulated with 11mM glucose and 10μM exendin-4. The results are plotted as individual insulin measurements. The difference between the groups was determined using the Mann-Whitney test. The data is pooled from 2 individual experiments.

### GLP1-R expression is decreased in the islets of patients with type 1 diabetes who retain endogenous C-peptide production

To assess whether our findings in mice had a potential relevance in human disease, we performed histological examination of human pancreas tissue from nPOD, the network for pancreas donors with diabetes (36). Included in the study were 6 healthy control donors, 7 donors with type 1 diabetes with retained C-peptide production (disease duration 0-22 years), 6 donors with type 2 diabetes (disease duration 2-25 years, none on insulin therapy, donor 6189 had received exenatide treatment), and 6 islet autoantibody-positive healthy donors. Type 2 diabetes specimens served as a control, as previous reports have demonstrated that hyperglycemia in itself can lead to a reduction in GLP-1R (37). Autoantibody positive individuals were included as these have been reported to have a higher risk of later developing type 1 diabetes (38).

To test whether we could detect differences in GLP-1R distribution, we stained sections from the donated pancreata for GLP-1R and insulin. Antibody specificity was tested beforehand, demonstrating that the GLP-1R antibody stained beta cells specifically and not alpha cells (Supplementary figure 7). Conditions for staining and analysis were kept consistent, allowing us to compare the intensity of GLP-1R and insulin staining in the islets in each section. We found that sections from people with type 1 diabetes all had significantly lower intensity for the GLP-1R staining (Fig. 7a, left panel) while the intensity for insulin staining, although showing a wider spread in sections from individuals with both type 1- and type 2 diabetes, was not significantly different (Fig. 7a, right panel). Representative islets for analysis are shown in Figure 7b, and larger fields are shown as tile scans in Figure 7c. Data from the individual specimens, and representative tile scans, are shown in Supplementary Figure 8.

**Figure 7.**
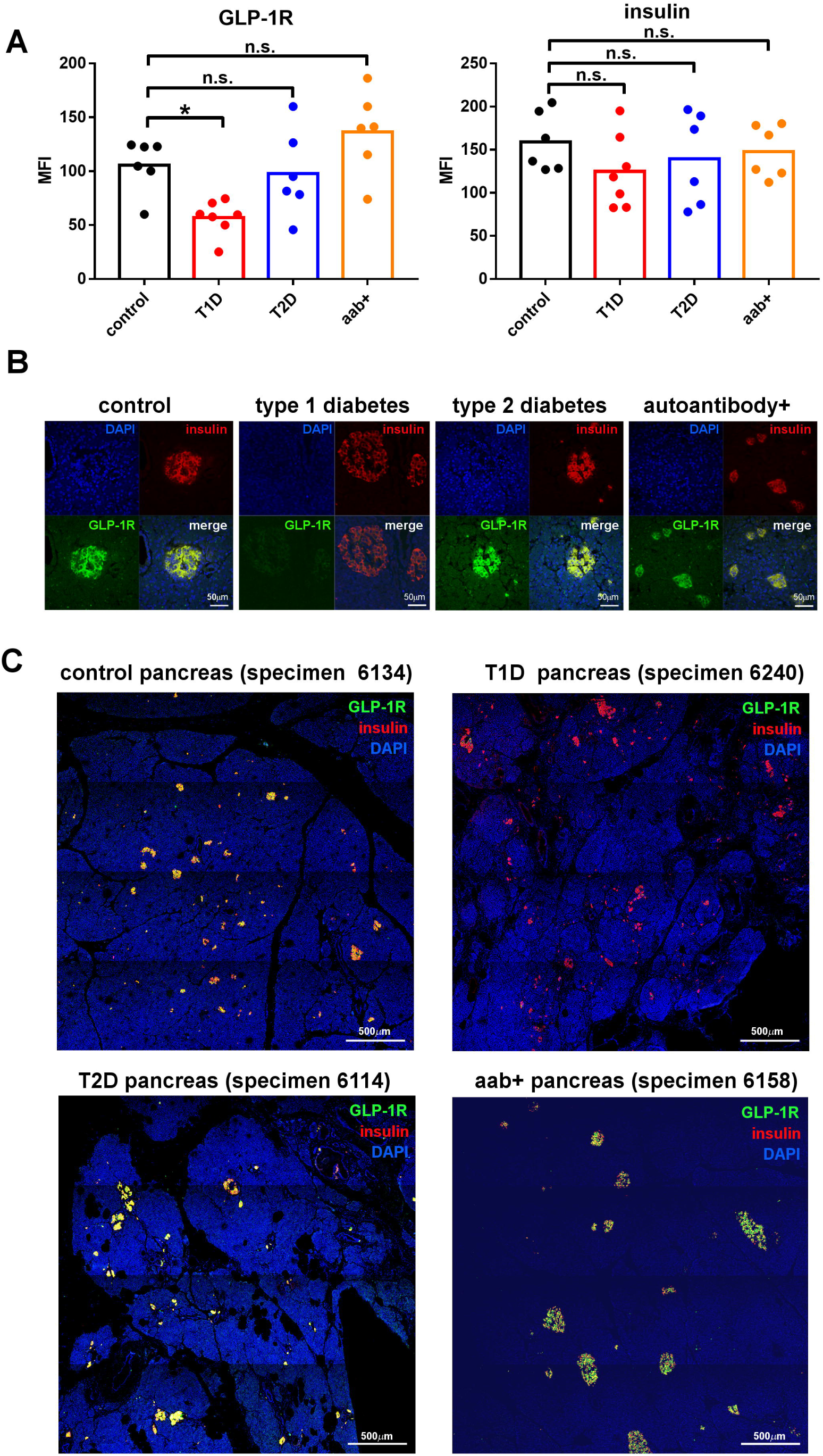
GLP1-R expression is decreased in the islets of patients with type 1 diabetes who retain endogenous c-peptide production. Paraffin embedded sections of pancreas samples from nPOD, from either healthy donors (control, black bars), people with type 1 diabetes (T1D, red bars), people with type 2 diabetes (T2D, blue bars), or healthy donors positive for islet autoantibodies (auto aab+, yellow bars) were stained with GLP-1R and insulin antibodies, and the mean intensity of staining for GLP-1R (A, left) and insulin (A, right) in at least 10 islets per section was plotted for each donor. Each dot represents the average of at least 10 islets from one donor. Differences between groups were assessed using ANOVA followed by a Dunnett’s post test for multiple comparisons. (B) Representative split images of analysed islets from control (left), type 1 diabetes (middle left), type 2 diabetes (middle, right) and islet autoantibody positive (right) donor samples. (C) Representative tile scans of sections show GLP-1R (green), insulin (red) and nuclear stain DAPI (blue).

## Discussion

Targeting the immune system to achieve immunological tolerance to islet antigens is an attractive approach to halting type 1 diabetes, but although delay of disease has been reported with treatments such as anti-CD3 antibody, it is not enough to achieve remission (39, 40). Some researchers have proposed that a combination of treatments targeting the immune response and the beta cells themselves may give better results (2, 41), and the already well-established drugs that target GLP-1R signalling, such as Byetta (exenatide, the generic drug name for exendin-4), stands out as a feasible and safe option. Attempts at combination treatment in type 1 diabetes with GLP-1R stimulation, although showing some efficacy, have also failed to achieve remission in mice (19) and humans (22), and the contribution of GLP-1R stimulation has been unclear. Studies investigating the effects of exenatide treatment in NOD mice reported a modest effect on diabetes incidence (18, 20). Our findings demonstrate that both diabetes susceptible NOD mice and people with type 1 diabetes express significantly lower levels of GLP-1R. These results may explain, at least in part, why exenatide, alone or in combination, has not achieved greater success in type 1 diabetes. Our conclusions are further corroborated by a study using labelled exendin-4 probes to image insulinomas via binding to the GLP-1R. The exendin-4 probe was unable to detect the pancreas in NOD mice but successfully detected the transplanted rat insulinoma cell line INS-1 (42).

There are indications that type 1 diabetes patients may have a dysregulated incretin effect, although this is difficult to assess given that the defining feature of the disease is the lack of insulin. A group of people diagnosed with type 1 diabetes with normal fasting blood glucose levels but impaired oral glucose tolerance test and presence of anti-islet autoantibodies displayed a significantly reduced incretin effect despite normal levels of GLP-1 in serum (43, 44). However, it is clear that some type 1 diabetes patients can respond to infusion of exogenous GLP-1, as a report from Vilsboll *et al.*(45) demonstrated that direct administration of GLP-1 iv to C-peptide positive volunteers with type 1 diabetes induced stimulation of insulin secretion, albeit with a large spread in responses (45).

The GLP-1R is central not just to beta cell function but also to beta cell stress resilience and regeneration. In NOD^*scid*^ mice, lower expression of the GLP-1R can be seen in the absence of any immune pathology in the islets, and this is seen even at an early age. This suggests that reduced levels of GLP-1R in NOD mice and the type 1 diabetes patients we studied might play a role in the initial events that lead to inappropriate presentation and priming of anti-islet immune responses in the first place, and contribute to the beta cell fragility observed in previous studies (6). To investigate whether a deficiency in GLP-1R precedes development of overt type 1 diabetes in humans, we requested nPOD samples from healthy donors with presence of islet autoantibodies. People with islet autoantibodies have a greater risk of developing type 1 diabetes (38), and we hypothesised that if GLP-1R deficiency precedes development of disease we would observe reduced levels of staining is the specimens from this donor group. However, these samples showed no decrease in the levels of GLP-1R, in fact four of the five highest recorded means of staining intensity were seen in this group. Thus our data, which we acknowledge is based on a small number of donor samples, does not indicate that downregulation of the GLP-1R precedes the formation of anti-islet immune responses in people.

Reduced expression of *glpr1* is a feature of immature beta cells, along with lower glucose stimulated secretion of insulin (46). We see no difference in *glpr1* message in the NOD mice, but there is a possibility that the reduced level of GLP-1R in the human donor samples we have assessed reflects higher turnover and an altered phenotype of the detected beta cells.

The GLP-1R gene does not map to any known type 1 diabetes susceptibility loci in either humans (3) nor mice (47), and there is no difference in the gene sequence between NOD mice and C57BL/6 mice (24). We did not detect any differences in mRNA levels between the mouse strains, but instead identified that the GLP-1R protein was continuously degraded in the NOD mouse beta cells. The GLP-1R is internalised upon ligand binding and recycles to the cell surface via endosomes (48). Signalling from the receptor continues after it has been internalised, and internalisation appears to instruct the strength and type of signalling resulting from ligand binding. However, ligands that induce less internalisation appear to achieve greater insulin release (48). When the receptor has been internalised, it can either be sorted back to the cell surface, or to the lysosomes where it is degraded (49). Our data suggest that the GLP-1R is present in the lysosomes in NOD mouse beta cells, and that downregulation of GLP-1R in NOD mice can be halted by inhibition of lysosomal degradation. It is interesting to note that *in vivo* treatment with chloroquine, a potent inhibitor of lysosomal degradation via its effects of acidification of the lysosomal compartment and thus on endosome fusion with the lysosome, leads to decreased incidence of diabetes in NOD mice (50) and to improved response in a glucose tolerance test in NOD mice (51). Chloroquine has many effects in an *in vivo* system and was used in those studies to probe the effects of TLR9, an intracellular toll like receptor that induces inflammation when it detects microbial DNA, but it is possible that the results also reflect an effect on GLP-1R that we see *in vitro* after treatment with chloroquine. A recent study identified several factors that contribute to turnover and signalling of the GLP-1R, such as clathrin, dynamin1 and Nedd4 (52), and future studies will reveal if any of these contribute to the increased degradation of the receptor in NOD mice. In contrast to the downregulation of the GLP-1R, we detected no deficiency in signalling from the GIPR in NOD mouse beta cells. This indicates that the two incretin receptors are differently regulated in NOD mouse beta cells and reveals an opportunity to pursue protection of the beta cell pool via simulation of this receptor as a therapeutic alternative. Treatment of type 1 diabetes using a combination of approaches targeting both the anti-islet immune response and the protection of the beta cell pool remains an attractive goal, and identification of pathways that lead to downregulation of the GLP-1R offers a new opportunity for therapy.

## Materials and methods

### Mice

NOD and NOD^*scid*^ mice used at the University of Cambridge and B6(Cg)-Tyr^c-2J^/J (JAX #000058) or white C57BL/6, NOD.Rag2^-/-^, NOD used at the Karolinska Institutet, Stockholm, Sweden, and white C57BL/6.Rag2^-/-^ mice used at the University of Lund (Sweden) were bred and maintained in a specific pathogen-free environment at the animal facilities and all procedures were approved by the ethical committee of Cambridge University, Karolinska Institutet, and Lund University Animal Care and Use, respectively. C57BL/6.*Rag*2^-/-^ mice were backcrossed to NOD mice for the generation of NOD.*Rag*2^-/-^ mice, as previously described (53) and white C57BL/6 were crossed to C57BL/6.Rag for the generation of white C57BL/6.Rag2^-/-^ mice for at least 5 generations. Blood glucose levels were measured using a Breeze2 blood glucose meter (Bayer). C57BL/6 mice were purchased from Charles River. The mice were used between 6-12 weeks of age and age and sex matched in all experiments. Mice were housed in IVC with free access to standard chow and water. This study was carried out in accordance with U.K. Home Office regulations.

### GLP-1 analogues

The GLP-1 peptide was purchased from Alta Biosciences(54) and the GIP from Tocris Biosciences. The labelled exendin-4 was either manufactured in the Department of Chemistry (TAMRA-Linker-HGEGTFTSDLSKQMEEEAVRLFIEWLKNGGPSSGAPPPS-NH2), University of Cambridge, or at Novo Nordisk, Copenhagen (Cy3-Linker-HGEGTFTSDLSKQMEEEAVRLFIEWLKNGGPSSGAPPPS-NH2). Exendin-4 was purchased from Sigma-Aldrich. Labelled exendin-9 (DLSKQMEEEAVRLFIEWLKNGGPSSGAPPPSC(Cy3)-NH2) was purchased from Cambridge Peptides.

For assessment of biological activity 400pmol of unlabelled exendin was injected iv, while up to 4nmol was injected i.v to assess binding of labelled probe as labelled probes have been demonstrated to have a significantly decreased affinity for the receptor(55).

### Section staining and microscopy

We were granted access to specimens from the network for organ donors with diabetes (nPOD)(36) (https://www.jdrfnpod.org/), from where we requested glass slides with sections of paraffin embedded pancreas from 7 healthy control donors (6007, 6029,6336, 6134, 6162, 6384 and 6339), 7 donors with type 1 diabetes with retained C-peptide production (6414, 6399, 6362, 6342, 6196, 6240 and 6405), 6 donors with type 2 diabetes (6273, 6108, 6114, 6139, 6189 and 6191) and 6 healthy donors with detected anti-islet autoantibodies (6123, 6080, 6314, 6090, 6147 and 6158, whereof all were positive for anti-GADA, and 6080 and 6158 were positive for anti-mIAA). Details about the donors such as disease duration, serum C-peptide and diabetes medication are included in supplementary table 1. The sections were dewaxed followed by antigen recovery, and then incubated with 1% H_2_O_2_ for 15 minutes to quench endogenous peroxidase. The sections were incubated with Mab 3F52 mouse anti-GLP-1R (DSHB, University of Iowa) (56) 1μg/ml for 2 h, followed by detection using the CSA-II kit (DAKO). Insulin was detected with guinea pig anti-insulin antibody (DAKO, Denmark) followed with secondary Alexa 547 anti-guinea pig (Invitrogen). Glucagon was detected with rabbit anti-glucagon antibody (Millipore) followed with secondary Alexa 547 anti-rabbit (Invitrogen). Nuclear staining was achieved with DAPI (Invitrogen). The sections were viewed with a LSM780 Confocal Microscope (Zeiss) and images were captured using the same settings for all sections acquired in the same session, with each session allocated an equal number of slides from each donor group to ensure congruity. The intensity of staining in individual islets was measured using ZEN software (Zeiss), from at least 10 randomly selected islets per section.

For staining of mouse GLP-1R in sections, Mab7F38 (57) (DSHB) was used with amplification using the CSA II kit (DAKO) according to the manufacturers instruction, and insulin and glucagon were detected with the same antibodies that were used for human sections as these are cross reactive.

### Inhibition of degradation

Freshly harvested islets were dispersed to single cell suspension using Enzyme Free Dissociation Buffer (Gibco), resuspended in DMEM medium (Sigma-Aldrich) without supplements and were plated on coverslips pre-coated with Poly-L-lysine (Sigma-Aldrich) in a damp chamber. After seeding at 37C°, cells were treated with Lysosome inhibitors (Chloroquine 50μg/ml and /or Bafilomycin 400nM, Sigma-Aldrich) or vehicle alone (DMSO) diluted in DMEM medium for 4h (37C°) and stained with LysoTracker DND-99 (Invitrogen). Cells were fixed in 4% PFA. For staining, cells were permeabilised with 0.5% Triton X-100 for 5mins in room temperature (RT) and blocked in 3% BSA/5% FCS/PBS for 1h. Staining was performed at RT for 1h by using Mab 7F38 mouse anti-GLP-1R (DSHB, University of Iowa) 1.5μg/ml, guinea pig anti Insulin (DAKO), rabbit anti-Calnexin (ENZO), rabbit anti-ZFPL1 (Atlas antibodies) and then with Alexa Fluor® conjugated-secondary antibodies (Invitrogen). Nuclei were stained with DAPI (Invitrogen). Coverslips were then mounted onto slides by using ProLong Diamond antifade mounting medium (Thermo Fisher Scientific) and allowed to dry over night before imaging. Images were acquired with LSM780 Confocal Microscope (Zeiss) and image settings were kept identical for all conditions. Zen software (Zeiss) was used to quantify the intensity of staining for at least 20 randomly selected islet cells per condition.

### Flowcytometry

Beta cells were prepared as described above and incubated in 1μM TAMRA-labelled exending-4 for 30 min at 37°C. The cells were then analysed on a Cytek DxP8 analyser, gated for a population enriched for beta cells (but not containing only beta cells) based on forward and side scatter, excluding doublets. The mean fluorescence intensity in the population positive for exendin-4, reported to be beta cells (33), was recorded.

### TaqMan PCR

RNA was extracted using the RNAEasy kit (Qiagen) and cDNA was generated as previously described (58). TaqMan PCR was performed using commercially available primer-probe sets for GLP-1R and GAPDH (Applied Biosystems) and analysed on an ABI Prism (Applied Biosystems) using SDS software, according to the manufacturer’s instructions. A standard curve was generated by performing a serial 1:10 dilution of cDNA from C57BL/6 islets, with the highest concentration given a value of 1, and used to translate the results into semiquantitative values. The amount of signal in each sample was calculated as the ratio between the target and the endogenous control, as indicated.

### IVIS imaging of tissues

Mice were injected with either Cy3 labelled exendin-4 (4nmol) or vehicle only iv, and one hour later the kidney, pancreas, liver, lungs heart, spleen and the whole gastrointestinal tract including the stomach were harvested and analysed in the IVIS imaging system (Perkin Elmer).

### Islet transplantation

#### Transplantation under kidney capsule

Islets were isolated as previously described (59) and transplanted under the kidney capsule as previously described (60). Islets were isolated from male NOD^*scid*^ or C57BL/6 mice (10-12 weeks old) and transplanted under the kidney capsule of either male NOD^*scid*^ or C57BL/6 recipients as indicated. After 10 days, the mice were given an injection of 4 nmol (approximately 2μg) of exendin4-Cy3 probe iv. One hour after the injection, the graft bearing kidney was harvested, fixed in 4% PFA, then embedded in OCT and the frozen sections were stained and analysed as described above.

#### Transplantation into the anterior chamber of the eye (ACE)

Mouse pancreatic islets were isolated and between 10 and 30 cultured islets were transplanted per eye, as previously described (31). Islets from white B6 mice were transplanted into eyes from 10-week-old female NOD.Rag2^-/-^ recipient mice (n=4) and non-inflamed islets from NOD.Rag2^-/-^ mice were transplanted into eyes from 7-week-old female white B6.Rag^-/-^ mice (n=3). At the time of transplantation, the recipient animal was anaesthetised using inhalation anaesthesia (isoflurane; Schering-Plough, Kenilworth, NJ, USA). To obtain postoperative analgesia, we administered buprenorphine (0.15mg/kg; RB Pharmaceuticals, Slough, UK) subcutaneously. Between 10 and 50 cultured islets were transplanted per eye as previously described (32). Injection of labelled exendin-4 was performed as described above.

### *In vivo* Two-Photon imaging and analysis

This method has been described previously (32). We used an upright Zeiss LSM 7 MP microscope system equipped with a tunable Ti:Sapphire laser (MaiTai, Spectra Physics) together with a long working distance water-dipping lens (Zeiss W Plan-Apochromat 20x/1.0 DIC M27 75mm). Viscotears (Novartis) was used as immersion liquid. For visualization of blood vessels, we intravenously injected 70-kDa-dextran-FITC (50 μl of 2.5 mg/ml; Invitrogen) followed by injection of inactive Exendin-4 (E9)-Cy3 (50μl, 0.5-2nmol in HBSS) via the tail vein. An injection of 0.5nmol of Exendin-4 (E9)-Cy3 sufficiently stained beta cells in B6 islet grafts within 30 min. We excited either FITC at 890nm or FITC and Cy3 at 1000 nm and collected emission light onto two non-descanned detectors using a dichroic mirror (RSP795) and emission filters (BP 500-550 and BP 565-610). For morphological characterization of a pancreatic islet graft reflection was imaged by exciting with 633 nm and measuring emission between 632 and 639 nm. Engrafted islets were imaged 10 weeks post transplantation for up to 40 min. Image sequences were recorded using typical imaging volumes of 512×512×10 for time series or 512×515×50 for whole islet acquisition, 0.639 × 0.639 × 3μm voxel size, z-spacing of 3μm or less.

### Cyclic AMP assay

Islets were isolated as previously described (59) and dispersed to single cell suspension using Enzyme Free Dissociation Buffer (Gibco). Islets were seeded in a 384-well Optiplate (PerkinElmer) (4000 cells per well) and stimulated with either GLP-1 or GIP for 30 minutes prior to measurement of cAMP accumulation using a LANCE® Ultra cAMP Detection Kit (PerkinElmer) in accordance with the manufacturer’s instructions and as described previously (61, 62). Results are presented as % of total stimulation with 10 μM forskolin (a direct activator of adenylyl cyclase).

### Insulin Assay

Islets from NOD^*scid*^ or C57BL/6 mice were handpicked and seeded in to a U-bottom 96 well plate (10 islets per well) containing either 11 mM glucose, but no Ca^2+^, alone or with exendin-4 (Sigma-Aldrich) added at the indicated concentrations. Extracellular Ca^2+^ is not required for GLP-1 induced glucose stimulated insulin secretion (34), and was excluded as Ca^2+^ added to the medium give a high background and variability. After 1h, supernatants were harvested and analysed for insulin content using the Mesoscale assay (MSD) in the Cambridge Biochemical Assay Laboratory, Addenbrooke’s Hospital.

### Statistics

We have used a paired Student’s t-test to compare before and after exendin-4 levels of blood glucose, and either Student’s t-test or Mann-Whitney tests to assess difference between groups in experiments with two groups as indicated in the figure legends. ANOVA followed by Dunnett’s post-test was used when more than two groups were assessed. Statistical significance is indicated as follows: *p≤0.05, **p≤0.01, ***p≤0.001, ****p≤0.0001, ns = non significant.

### Study approval

All animal experiments and procedures were approved by the were approved by the Animal Welfare and Ethical Review Body of Cambridge University, Karolinska Institutet, and Lund University Animal Care and Use, respectively. The study of human tissue samples provided from nPOD was carried out with the approval of the Human Biology Research Ethics Committee of the School of Biological Sciences, University of Cambridge.

## Supporting information

graphical abstract

## Acknowledgements

We thank the BSU staff and Yvonne K. Sawyer for technical support. We wish to thank Dr Alessandra De Riva, Department of Medicine, University of Cambridge, for the gift of NOD mice from her colony. We thank Pia Gottrup Mortensen and Dr Charles Pyke (Novo Nordisk, Copenhagen) for advice on staining protocols for Mab3F52. We thank the Cambridge Biochemical Assay Laboratory (CBAL) for technical assistance.

