## Supplementary figures for "GLP-1R is downregulated in beta cells of NOD mice and T1D patients"

S1

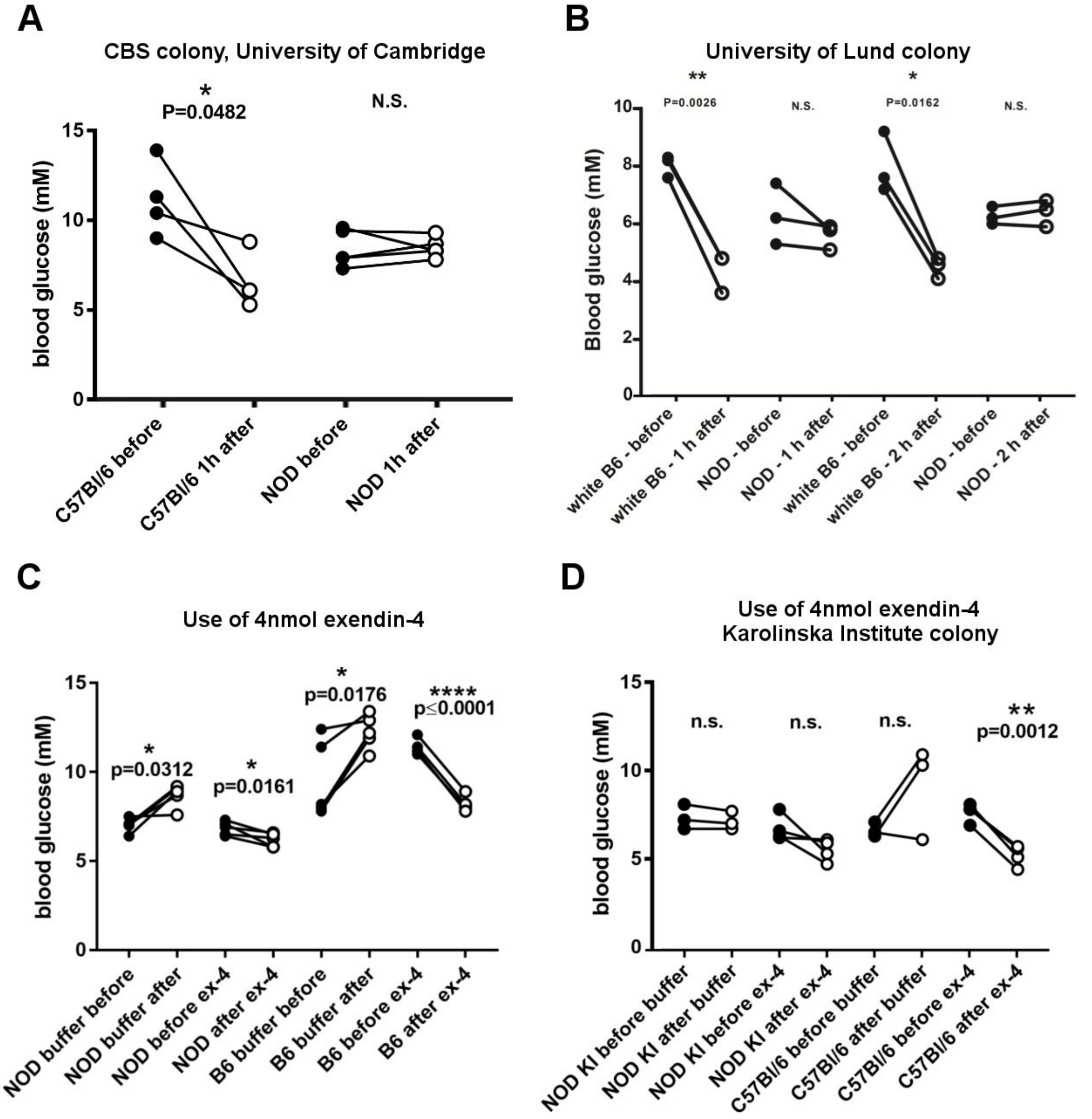


**S1. NOD mice from other colonies are also hyporesponsive to exendin-4 injection, and increasing the dose 10-fold can induce an attenuated response.** (A, B) Measurements of blood glucose before and after injection of 400 pmol exendin-4, or vehicle alone, in mice from the colonies kept at the Addenbrooke’s Hospital Campus, University of Cambridge (A), and at the University of Lund (B). (C, D) Measurement of blood glucose before and after injection of 4 nmol exendin-4, or vehicle alone, in mice from the colonies kept at the Department of pathology, University of Cambridge (C), and at the Karolinska Institute (D). Differences were measured using a paired t-test comparing before and after blood glucose levels. Each dot represents one mouse.

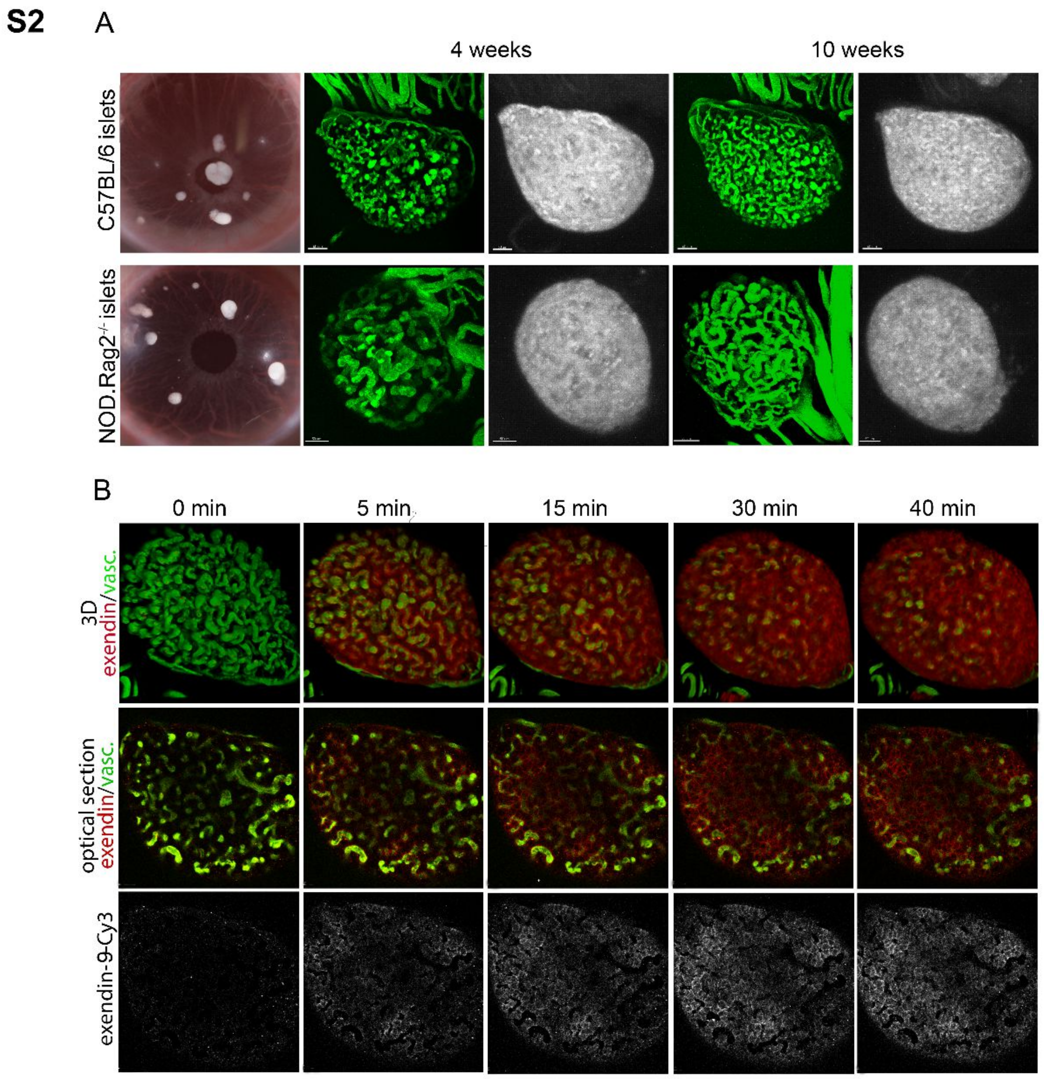


**S2. Longitudinal imaging of ACE-transplanted C57BL/6 or NOD.Rag2-/- islets.** (A) Islets were cross-transplanted into the ACE of 10-week-old immunodeficient NOD.Rag2-/- or albino C57BL/6.Rag2-/- recipients (n=3 mice each) and individual islets were repeatedly imaged by 2-Photon microscopy at the indicated time points after transplantation. The column to the left displays photographs of the recipient mouse eye transplanted with islets engrafted on the iris. The increasing vascular density over time was visualized by intravenously administered dextran-FITC, 70 kDa, at the indicated time points, displayed as maximum projections in green. Islet morphology is displayed as maximum projections of islet reflection. (B) Ex4-Cy3 mean probing intensity increases progressively in C57BL/6 islets, respectively, reaching maximum staining intensity at approx. 30 min post injection. NOD.Rag2-/- recipient mice (n=3) transplanted with C57BL/6 islets were injected with 2nmol Ex4-Cy3 probe (shown in red) at 4 weeks post transplantation and individual islets were imaged repeatedly at indicated time points prior (0 min) or post injection. A representative imaging session of one individual islet is displayed as three-dimensional renderings of whole islet stacks (top row) or optical sections (middle and bottom row) at indicated timepoints.

**S3**

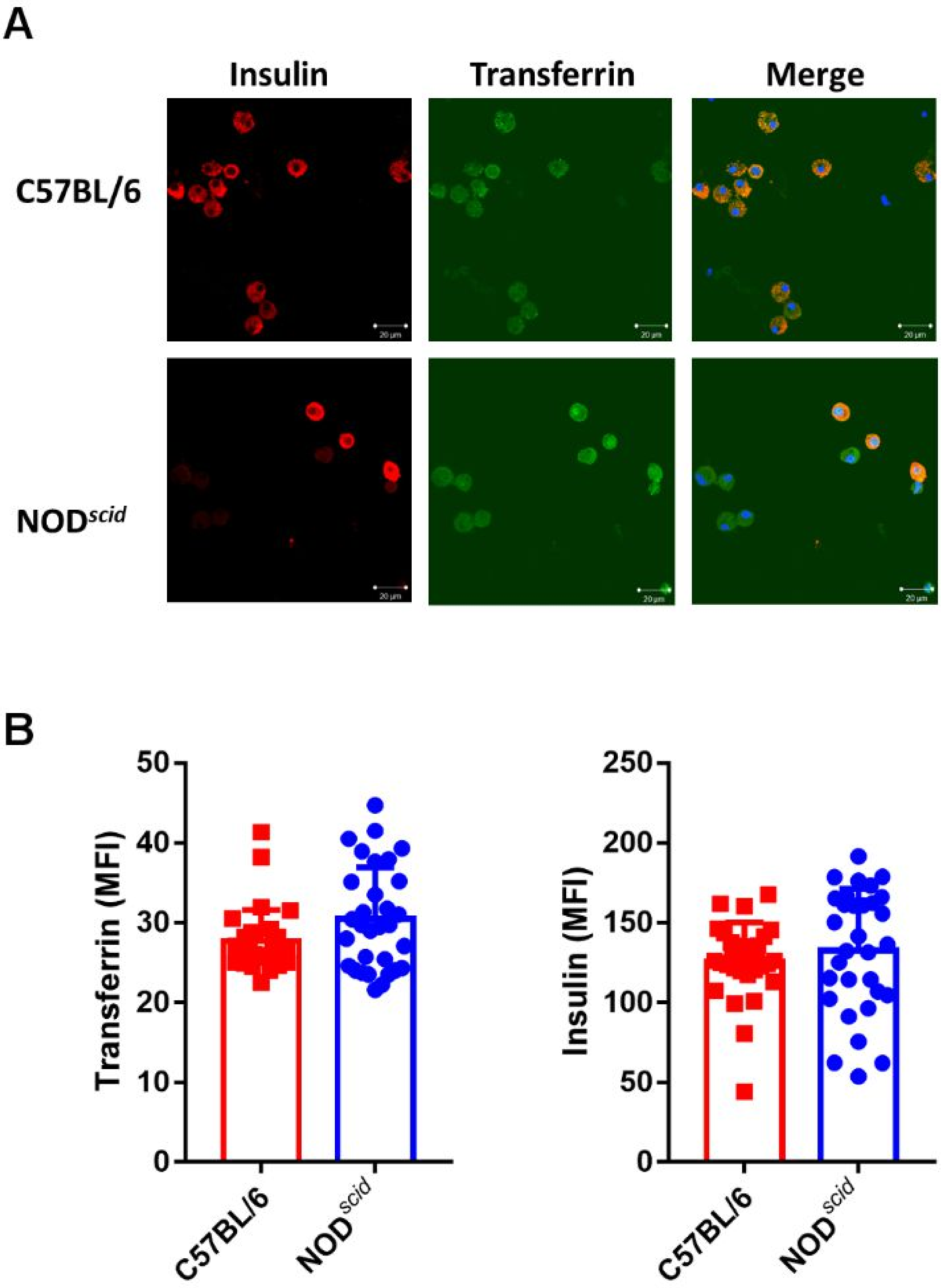


**S3. Expression of transferrin receptor is not lower in NOD*scid* beta cells than in C57BL/6 beta cells.** (A) Co-staining with Insulin (red) and Transferrin Receptor (green) in C57BL/6 (top panels) or NOD*scid* beta cells (lower panels). Nuclear stain DAPI is shown in blue. (B) Graphs plotting the recorded mean intensities for transferring (left) and insulin (right). Each dot represents one cell and testing for differences between groups was performed with Mann-Whitney test.

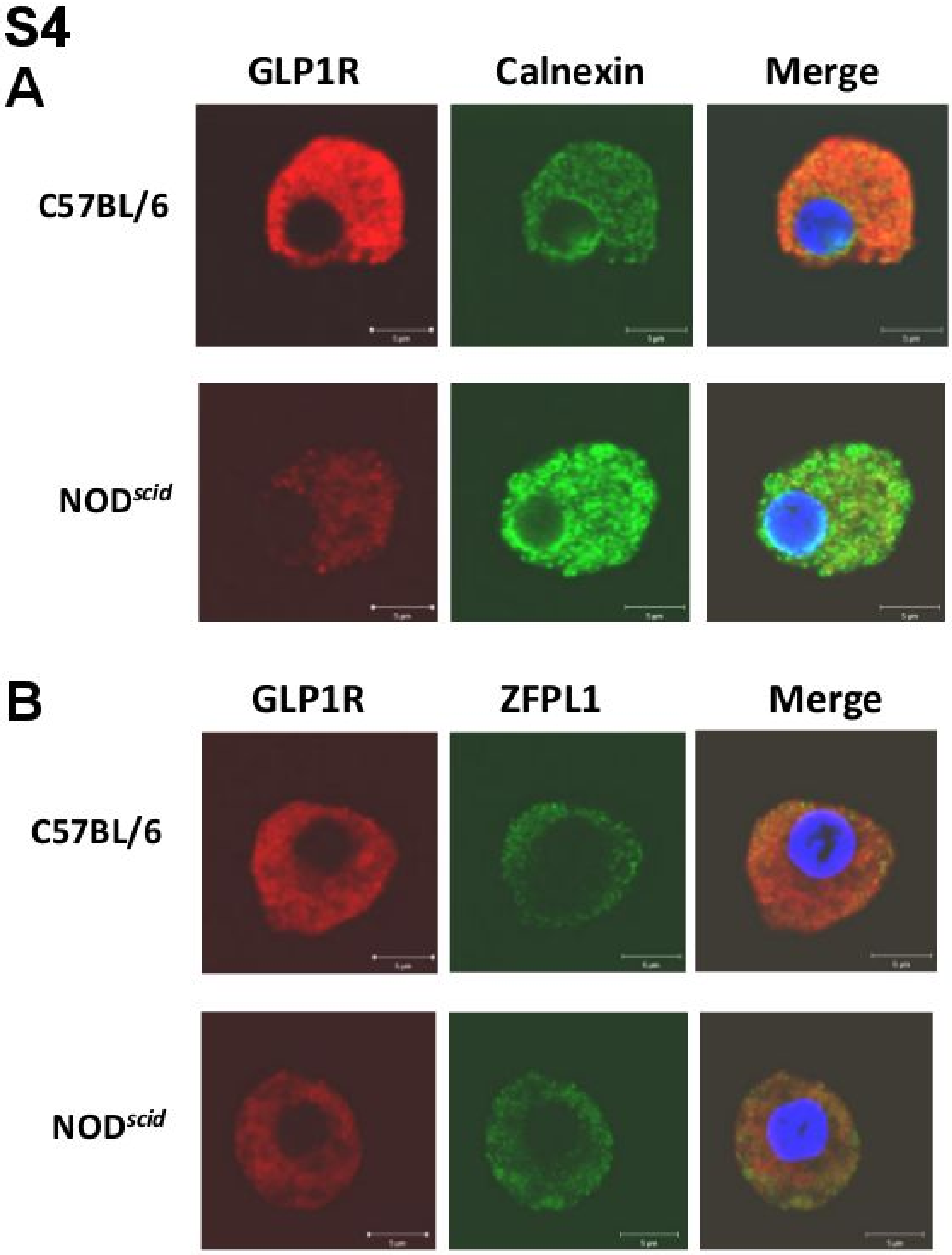


**S4. GLP-1R is not preferentially retained in the endoplasmic reticulum or the Golgi in NOD*scid* beta cells.** Co-staining with GLP-1R (red) and ER marker Calnexin (green) (A) or Golgi marker ZFPL1 (green) (B) in C57BL/6 or NOD*scid* beta cells. Nuclear stain DAPI is shown in blue.

S5

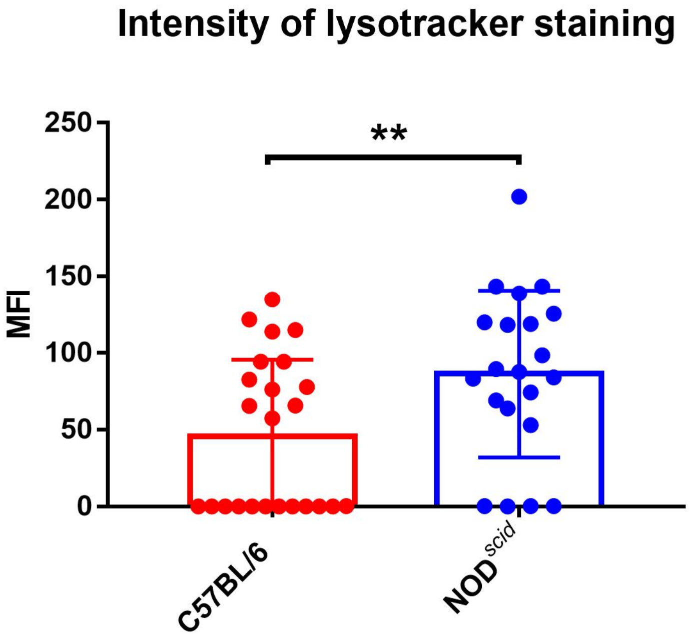


**S5. NOD*scid* beta cells have higher levels of lysotracker staining, as measured by mean fluorescence intensity in beta cells.** Each dot represents one cell. Differences were evaluated using a Mann-Whitney test.

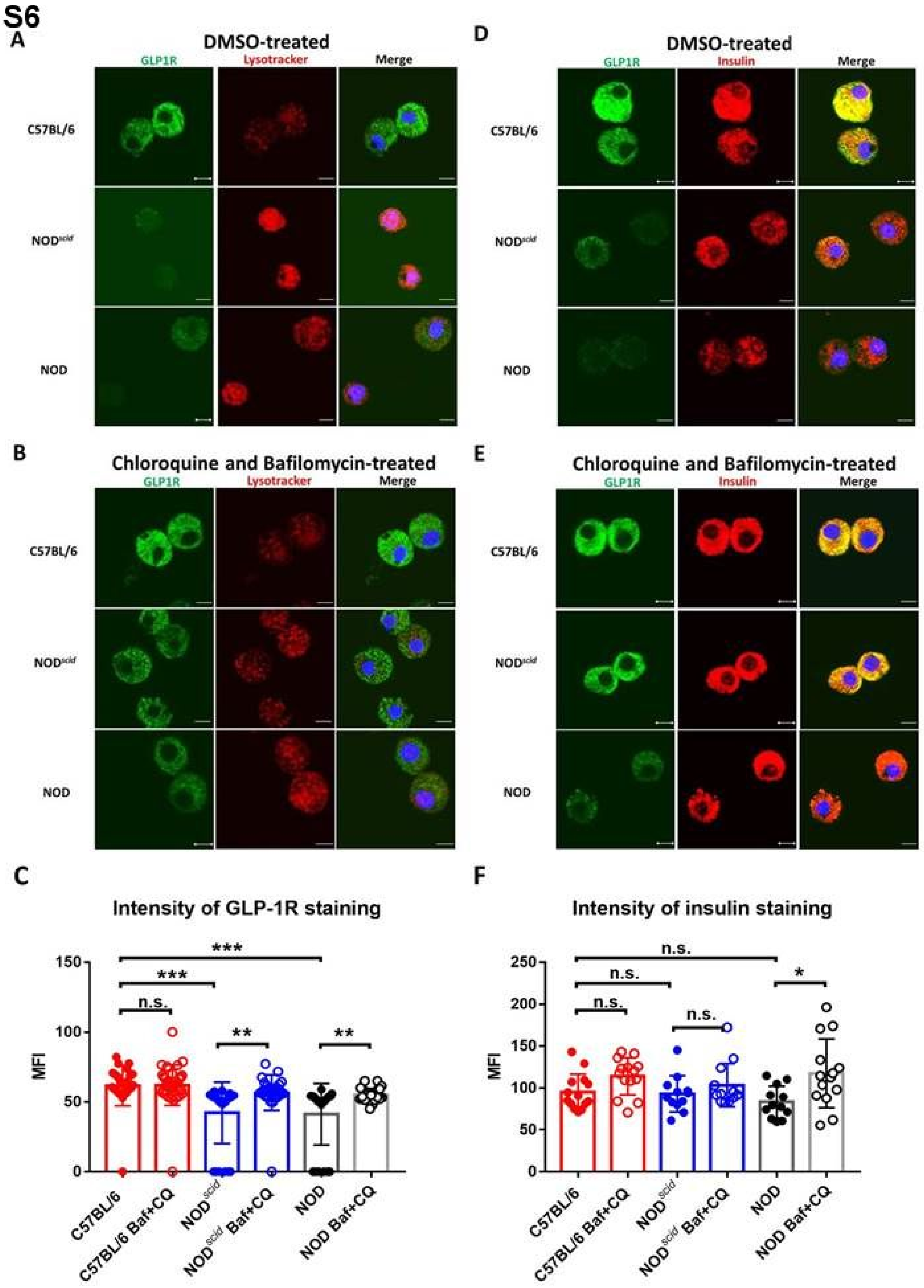


**S6. Insulin content in C57BL/6, NOD and NOD$^{scid}$ beta cells is not reduced by incubation with chloroquine and bafilomycin.** Beta cells from C57BL/6, NOD and NOD$^{scid}$ mice were incubated with vehicle (DMSO) alone (A, D), or a combination of chloroquine and bafilomycin for 4 h (B, E), after which the intensity of GLP-1R staining (A, B) and insulin staining (D, E) was measured by microscopy. (C, F) Graphs plotting the recorded mean intensities for GLP-1R (C) and insulin (F). Each dot represents one cell and testing for differences between groups was performed with ANOVA followed by Dunnett's test for multiple comparisons.

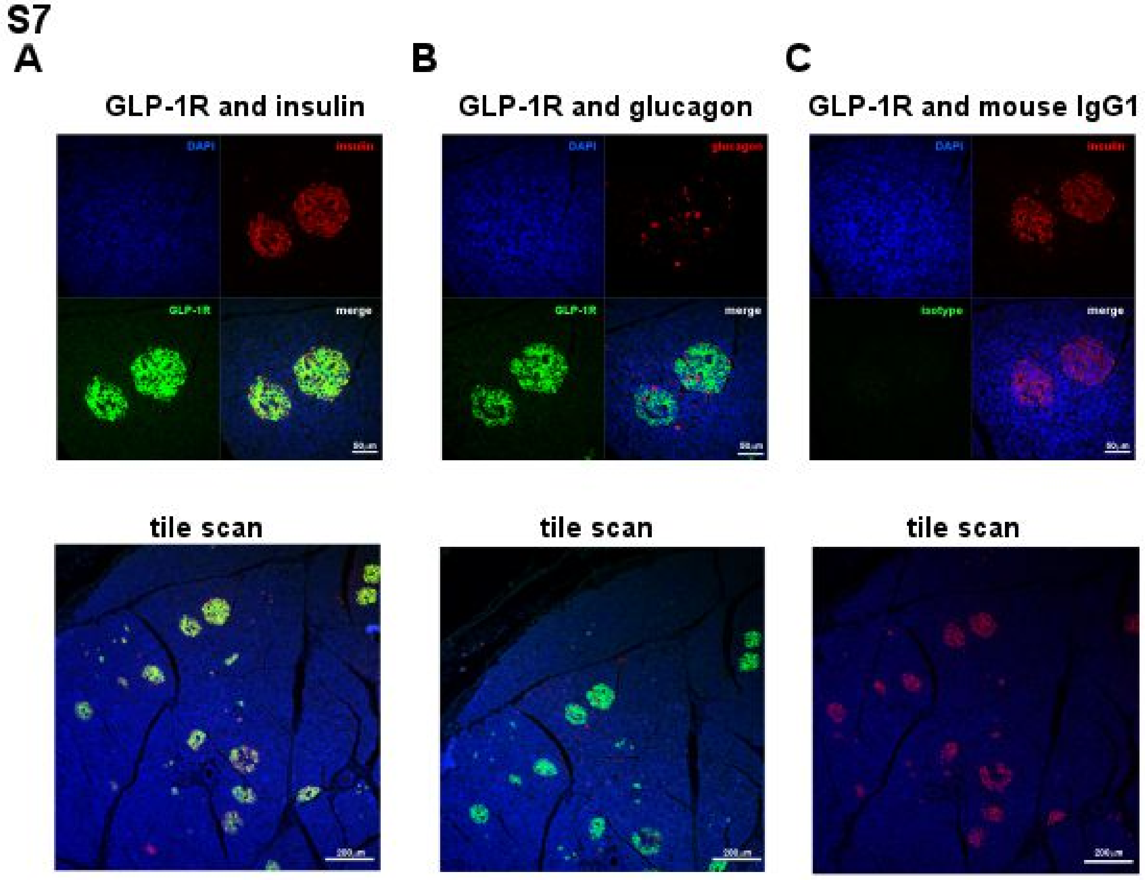


**S7. GLP-1R staining co-localises with insulin, but not glucagon staining in islets from healthy human pancreas.** Healthy human pancreas tissue (nPOD, specimen 6011) was stained for GLP-1R (green) and insulin (red, A) or glucagon (red, B). Mouse IgG1 antibody was used as an isotype control to demonstrate binding specificity (C). Top panels show split images of all channels, while the bottom panels show tile scans of a larger area of the stained sections.

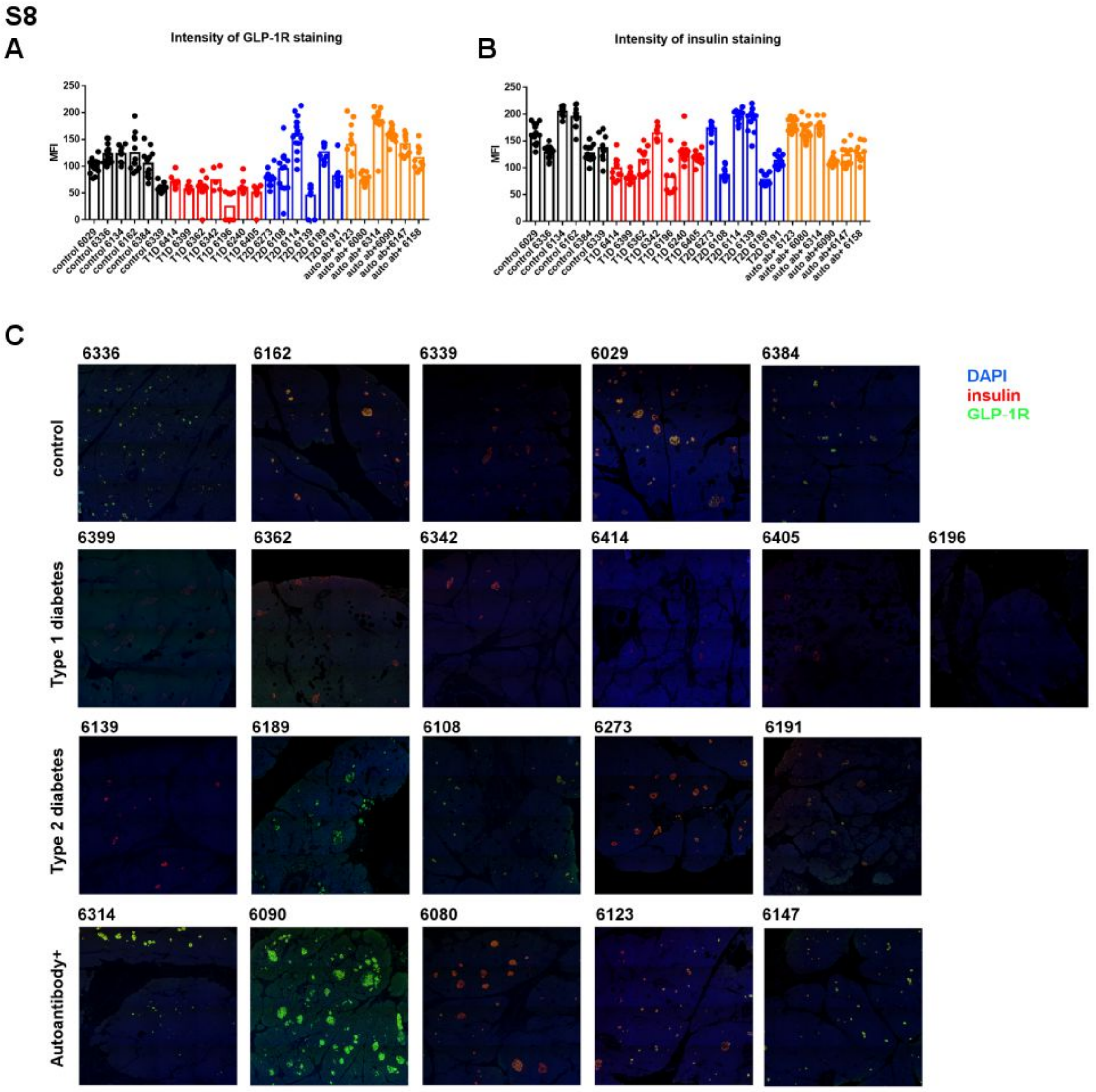


**S8. Islets in pancreas tissue from people with type 1 diabetes have lower levels of GLP-1R.** Paraffin embedded sections of pancreas samples from nPOD, from either healthy donors (control, black bars), people with type 1 diabetes (T1D, red bars), people with type 2 diabetes (T2D, blue bars), or healthy donors positive for islet autoantibodies (auto aab+, yellow bars) were stained with GLP-1R and insulin antibodies, and the mean intensity of staining for GLP-1R (A) and insulin (B) in at least 10 islets per section was plotted for each donor. Each dot represents one islet. Tile scans of each section show GLP-1R (green), insulin (red) and nuclear stain DAPI (blue) (C).

### Supplementary table 1

| nPOD CaseID | Donor Type | Age (yrs) | Diabetes duration (yrs) | C-peptide | Peak glucose (mg/dl) | HbA1c | Diabetes medication | Sex | Race | BMI (kg/m2) |
|---|---|---|---|---|---|---|---|---|---|---|
| 6342 | T1D | 14.0 | 2 | 0.26 | 889 | 9.2 | Insulin | Female | Caucasian | 24.3 |
| 6362 | | 24.9 | 0 | 0.38 | 435 | 10 | Insulin IV drip on admission | Male | Caucasian | 28.5 |
| 6367 | | 24.0 | 2 | 0.39 | 512 | 8.8 | Novalog, Lantus | Male | Caucasian | 25.7 |
| 6399 | | 17.4 | 0 | 1.41 | 867 | 10.4 | None | Male | Caucasian | 32.0 |
| 6405 | | 29.1 | 0.6 | 1.84 | 188 | 7 | Insulin | Female | Hispanic/Latinx | 42.5 |
| 6414 | | 23.1 | 0.43 | 0.16 | 690 | 14 | Lantus and Novalog | Male | African Am | 28.4 |
| 6196 | | 26.5 | 15 | 0.48 | 860 | | Insulin | Female | African Am | |
| 6080 | Autoab Pos | 69.2 | | 1.84 | 226 | | | Female | Caucasian | 21.3 |
| 6090 | | 2.2 | | 5.34 | 402 | | | Male | Hispanic/Latinx | 18.8 |
| 6123 | | 23.2 | | 2.01 | 267 | 5.4000 | | Female | Caucasian | 17.6 |
| 6147 | | 23.8 | | 3.19 | 287 | 5.2000 | | Female | Caucasian | 32.9 |
| 6158 | | 40.3 | | 0.51 | 449 | 5.6000 | | Male | Caucasian | 29.7 |
| 6314 | | 21.0 | | 1.49 | 207 | | | Male | Caucasian | 23.8 |
| 6029 | No diabetes | 24.0 | | | | | | Female | Hispanic/Latinx | 22.6 |
| 6134 | | 26.7 | | 3.59 | | | | Male | Caucasian | 20.1 |
| 6162 | | 22.7 | | 7.61 | 312 | | | Male | African Am | 28.9 |
| 6336 | | 14.3 | | 7.87 | 294 | 5.2 | | Female | Caucasian | 28.9 |
| 6339 | | 23.3 | | 10.56 | 214 | 5.3 | | Male | Caucasian | 25.0 |
| 6384 | | 17.0 | | 0.7 | 305 | 4.8 | | Male | Caucasian | 18.2 |
| 6108 | T2D | 57.9 | 2 | 1.25 | 288 | | Metformin | Male | Asian | 30.4 |
| 6114 | | 42.8 | 2 | 0.58 | 400 | 7.8 | Metformin | Male | Caucasian | 31.0 |
| 6139 | | 37.2 | 1.5 | 0.6 | 513 | | None per chart | Female | Hispanic/Latinx | 45.4 |
| 6189 | | 48.5 | 26 | 1.85 | 198 | | Exenatide, Metformin, Glipizide, Insulin | Female | Caucasian | 36.1 |
| 6191 | | 62.7 | 10 | 6.14 | 265 | 6 | Metformin | Female | Caucasian | 19.9 |
| 6273 | | 45.0 | 2 | 3.17 | 286 | | Metformin | Female | African Am | 39.1 |

Table 1. Details of the nPOD specimens used in this study.
