## Supplementary figures and images for "GLP-1R is downregulated in beta cells of NOD mice and T1D patients"

### graphical abstract

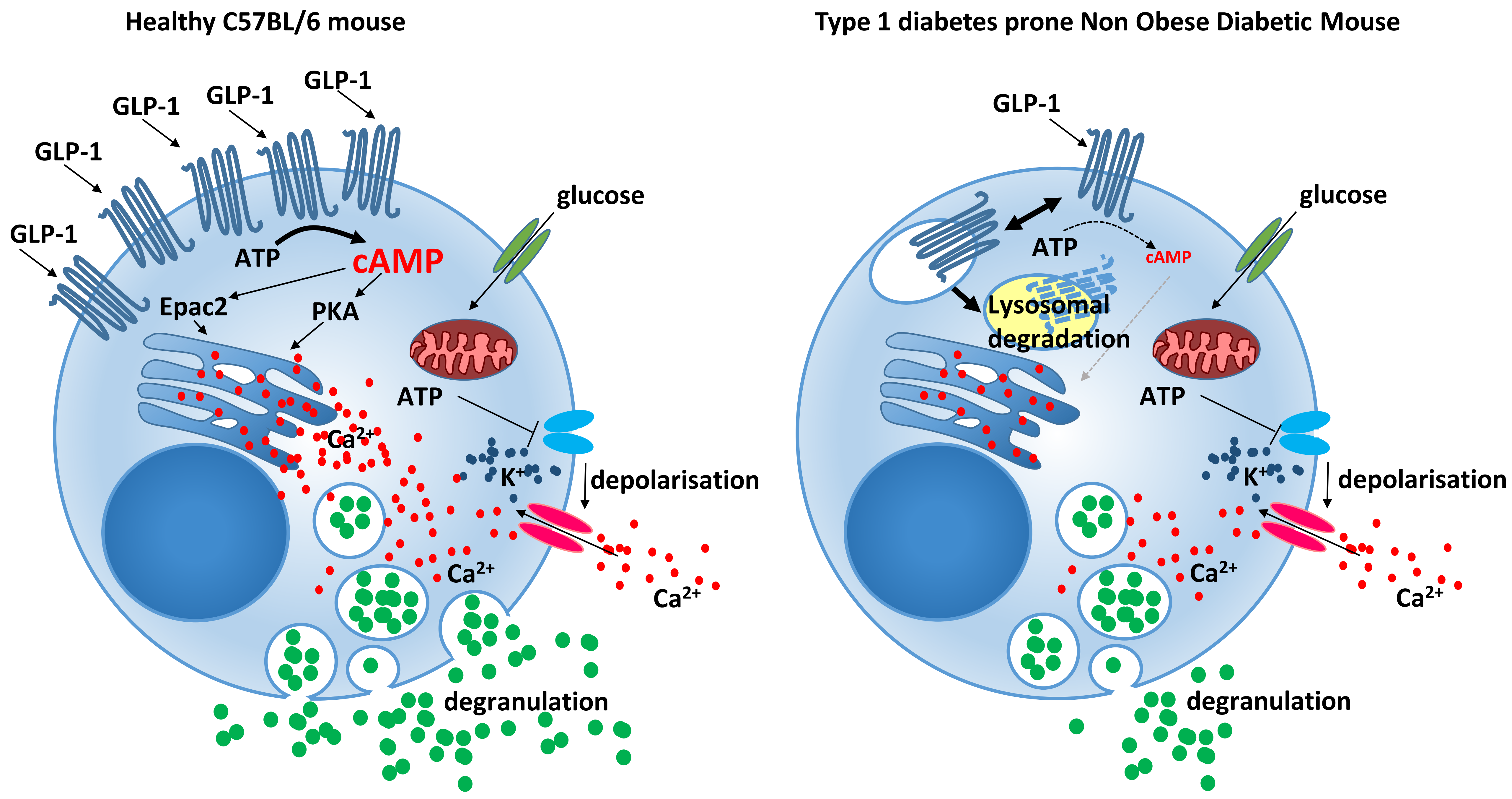
